# The RNA virome of intertidal kelp in New Zealand reveals novel viral lineages associated with unhealthy kelp

**DOI:** 10.64898/2026.02.05.704085

**Authors:** Lily E. M. Harvey, Stephanie J. Waller, Rebecca K. French, Jemma L. Geoghegan, Ceridwen I. Fraser

## Abstract

Pathogens can have major impacts on ecosystem function, especially when they infect foundational taxa such as habitat-forming kelp. Understanding which potentially pathogenic microbes are in an ecosystem is increasingly important as environmental change intensifies ecosystem imbalances, yet few studies have examined which viruses are associated with key habitat-forming macroalgae. Here, we screened multiple populations of intertidal / shallow-subtidal macroalgae in New Zealand for viruses. Samples were collected from three southern bull kelp species: *Durvillaea antarctica*, *Durvillaea poha* and *Durvillaea willana*. Total RNA was extracted and sequenced from both obviously ‘unhealthy’ kelp tissue and apparently ‘healthy’ tissue. We identified several novel viruses belonging to the families *Mitoviridae*, *Narnaviridae*, *Partitiviridae*, *Totiviridae*, and the recently proposed Ormycovirus group. Viral abundance and diversity were notably higher at Lawyer’s Head (the site closest to a city) compared to other sites, especially in unhealthy kelp samples. The elevated viral presence at Lawyer’s Head may indicate that human impacts, such as pollution, are influencing viral prevalence. This study reveals that coastal macroalgae in New Zealand host diverse viral lineages, some likely pathogenic, and hints at the potential influence of anthropogenic pollution on marine virome composition.

## Introduction

Marine microorganisms, which include bacteria, fungi, viruses, and protists, are highly abundant and diverse, driving large-scale biogeochemical processes and productivity across global ocean systems due to their extremely high biomass and ecological influence (Breitbart *et al*., 2008; Mojica & Brussaard, 2014; Ducklow, 2008). While most microorganisms are not pathogenic, those that are can significantly affect ecosystems by influencing host population dynamics, which can lead to cascading effects on broader ecosystem processes (Tompkins *et al*., 2011). Understanding pathogen-host interactions is essential for understanding ecosystem functions, species interactions, and services, especially as climate change may alter pathogen ranges and increase host vulnerability to disease (Mabey *et al*., 2021; Tylianakis *et al*., 2008; Hing *et al*., 2016).

Viruses dominate marine ecosystems in terms of sheer number, and greatly influence energy flow, nutrient cycling, diversity, primary productivity, and food web dynamics in the ocean through the infection and mortality of marine microorganisms (Breitbart *et al*., 2008; Mojica & Brussaard, 2014). Viral-host relationships can vary from pathogenic or antagonistic, where the virus damages the host, to commensal, where the virus has no detectable impact on the host, and mutualistic, where the virus benefits the host (Roossinck & Bazán, 2017). Pathogenic viruses have far-reaching consequences in the ocean, causing mortality and disease in a range of organisms including marine mammals, fish, molluscs, crustaceans and macroalgae (Lang *et al*., 2009; Schroeder & Mckeown, 2021). In macroalgae, viruses have been detected across diverse lineages, with evidence that they can influence host physiology and disease expression (Lachnit *et al*., 2016; van der Loos *et al*., 2023; Ruiz Martinez *et al*., 2023; Benites & Alves-Lima, 2022). Several studies suggest a potential role of viruses as pathogens in kelps, including associations between viral sequences and symptomatic kelp tissue (Beattie *et al*., 2018; McKeown *et al*., 2018; Zhang *et al*., 2022; Zhang *et al*., 2024). These findings indicate that viruses are both widespread in seaweeds and may play roles in macroalgal health and disease.

*Durvillaea*, commonly known as southern bull kelp or rimurapa in Aotearoa (New Zealand), is a genus of large brown macroalgae distributed across the Southern Hemisphere, with populations along the coasts of New Zealand, Australia, South America, and the sub-Antarctic islands (Blake *et al*., 2017; Velásquez *et al*., 2020). Three species occur on mainland New Zealand: two buoyant species: *D. antarctica* and *D. poha* and one non-buoyant *D. willana* (Fraser *et al*., 2020; Velásquez *et al*., 2020). *Durvillaea* is harvested for use as a food source in both Chile and Australia. In New Zealand, it holds cultural importance for Māori (the indigenous population of New Zealand), who traditionally use the kelp to create pōhā, bags that are used to store and transport tītī (muttonbirds) (Velásquez *et al*., 2020). Due to stressors such as marine heatwaves caused by climate change, *Durvillaea spp.* are threatened by range reduction, mortality, invasive species, and reduced fitness (Thomsen *et al*., 2019). It is therefore especially important to understand what pathogens are present in *Durvillaea* populations and the host-pathogen interactions for monitoring and conservation in the future.

The loss of keystone macroalgal species, such as bull kelp, has been shown to result in ecological shifts, including the proliferation of invasive species and declines in native species, highlighting the potential for bottom-up effects in marine ecosystems (Thomsen *et al*., 2019). Pathogens can impact macroalgal populations by reducing individual density, impairing reproductive potential, causing widespread mortality, and weakening blades and holdfasts, which reduces structural integrity, flexibility, and resistance to hydrostatic pressures (Neuhauser *et al*., 2011; Mabey *et al*., 2021). Widespread mortality and thinning can make macroalgal populations more susceptible to invasion by other algal species, which can lead to phase shifts from kelp forests to altered ecosystems over time (Reeves *et al*., 2022).

Despite the ecological and cultural importance of *Durvillaea*, pathogens associated with this kelp species are understudied. To date, no viruses have been identified infecting *Durvillaea*, and studies of viruses in macroalgae overall are limited (Schroeder & Mckeown, 2021). However, previous surveys of macroalgal viromes indicate that viruses are common in seaweed microbiomes, and in some cases have been proposed as candidate pathogens in kelps (Beattie *et al*., 2018; McKeown *et al*., 2018). These emerging insights highlight a significant gap in knowledge of viral diversity and function in *Durvillaea* specifically. Only three pathogen species have been described in *Durvillaea*: phytomyxean protistan parasites *Maullinia braseltonii* and *Maullinia ectocarpii*, and the algal endophyte *Herpodiscus durvillaeae* (Murúa *et al*., 2024; Blake *et al*., 2017; South, 1974).

Populations of *Durvillaea* occur across the Southern Hemisphere and are separated by large geographical distances. Buoyant species of *Durvillaea* have gas-filled honeycomb structures inside blades enabling them to float and travel extensive distances (Fraser *et al*., 2020). Solid-bladed *Durvillaea* could also potentially be dispersed with buoyant *Durvillaea* or other buoyant macroalgae such as *Macrocystis* (Fraser *et al*., 2020; Kelly *et al*., 2021). Kelp rafts could facilitate the introduction of hitchhiking invertebrates (e.g., Fraser *et al*., 2011) or microorganisms (Pearman *et al*., 2024) to new locations. Indeed, two non-viral pathogens of *Durvillaea*, *M. braseltonii* and *H. durvillaeae*, are both inferred to have crossed the Southern Ocean recently, with their buoyant hosts (Fraser & Waters, 2013; Mabey *et al*., 2021).

Climate change may alter the spread and host-pathogen dynamics in the marine environment. Climate change may cause physiological stress in hosts due to changes in abiotic factors, resulting in decreased fitness and increased susceptibility to infectious disease (Hing *et al*., 2016). Changing abiotic factors such as temperature may facilitate pathogens as their thermal tolerance can be greater than their hosts (Byers, 2021; Mojica & Brussaard, 2014). Climate change is predicted to continue altering oceanic circulation patterns, intensifying, weakening, or altering paths of fronts and currents, potentially resulting in changes in the dispersal of marine organisms (Wilson *et al*., 2016). Changes in the dispersal of marine life may alter species introductions and range, including the microorganisms that infect them. Pathogen propagule pressure could increase with intensifying and shifting currents, leading to increased infection rates and introductions of new diseases.

Understanding the pathogens present in different marine species and populations is crucial for tracking and mitigating new introductions that could disrupt these ecosystems, especially in the context of climate change. We here aimed to screen apparently unhealthy (and sympatric healthy) kelp tissue for viruses. Based on the wide range of disease-like symptoms observed in the field we hypothesised that *Durvillaea* hosts diverse viruses, some of which are likely to be pathogenic. We also hypothesised that viral abundance and virus family richness would differ between apparently unhealthy versus ‘healthy’ tissue, and among sites, due to environmental influences.

## Materials & Methods

### Kelp Sample Collection

Samples were collected from three sites in Otago and Southland, southern New Zealand, between March and May 2024 (Tautuku Peninsula in The Catlins, Brighton Beach just south of the city of Dunedin, and Lawyer’s Head in Dunedin) (Fig. 1 & Supplementary Table 1). Samples were collected from Brighton Beach on two separate occasions. Southern bull kelp individuals were assessed for visual signs of disease such as lesions, discolouration, deformities and necrosis, and biopsy samples of both seemingly unhealthy and seemingly healthy individuals were taken (Fig. 2). For ‘unhealthy’ tissue, photos were taken before cutting any tissue and descriptions of symptoms were noted (Supplementary Table 2). Pieces < 1 cm^2^ were cut from the middle of unhealthy areas using clean scissors and placed in tubes containing 800 μl of DNA/RNA Shield (Zymo Research: ZR) to preserve nucleic acids without freezing samples during collection and transport. Scissors were cleaned by a seawater rinse in the field. The same method was used to cut healthy tissue, from areas with no visual deformities, discolouration or lack of honeycomb in buoyant species. *Durvillaea antartica* was the most challenging species to sample due to inhabiting areas with much greater wave exposure than *D. willana* and *D. poha*, so fewer samples were collected (Supplementary Table 1).

**Fig. 1.**
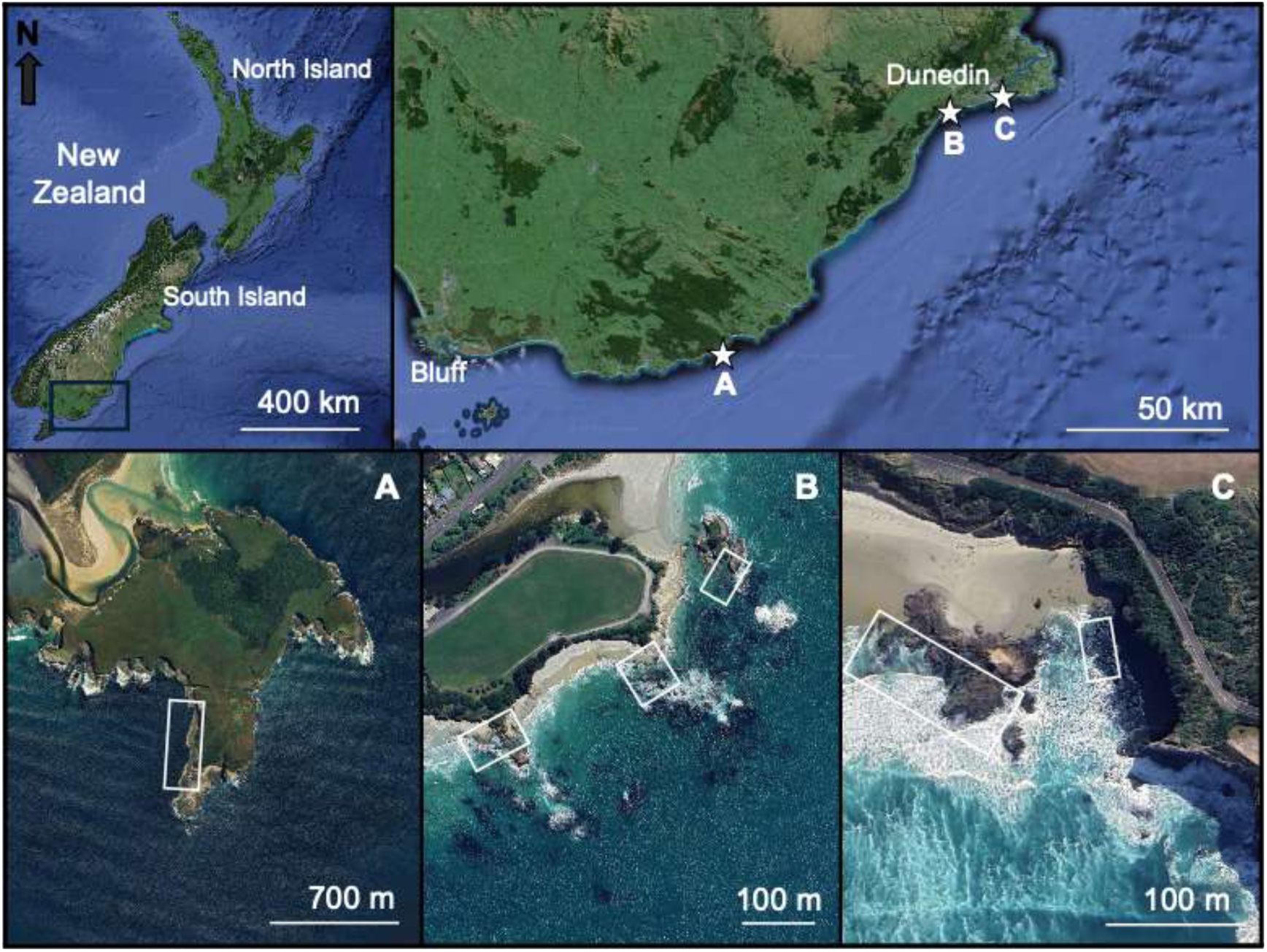
Maps showing the location of sites across Otago, New Zealand. White stars show the location of each site. A is Tautuku Peninsula, B is Brighton Beach, and C is Lawyers Head. White boxes show areas where samples were collected. Satellite images were sourced from Google Earth.

**Fig. 2.**
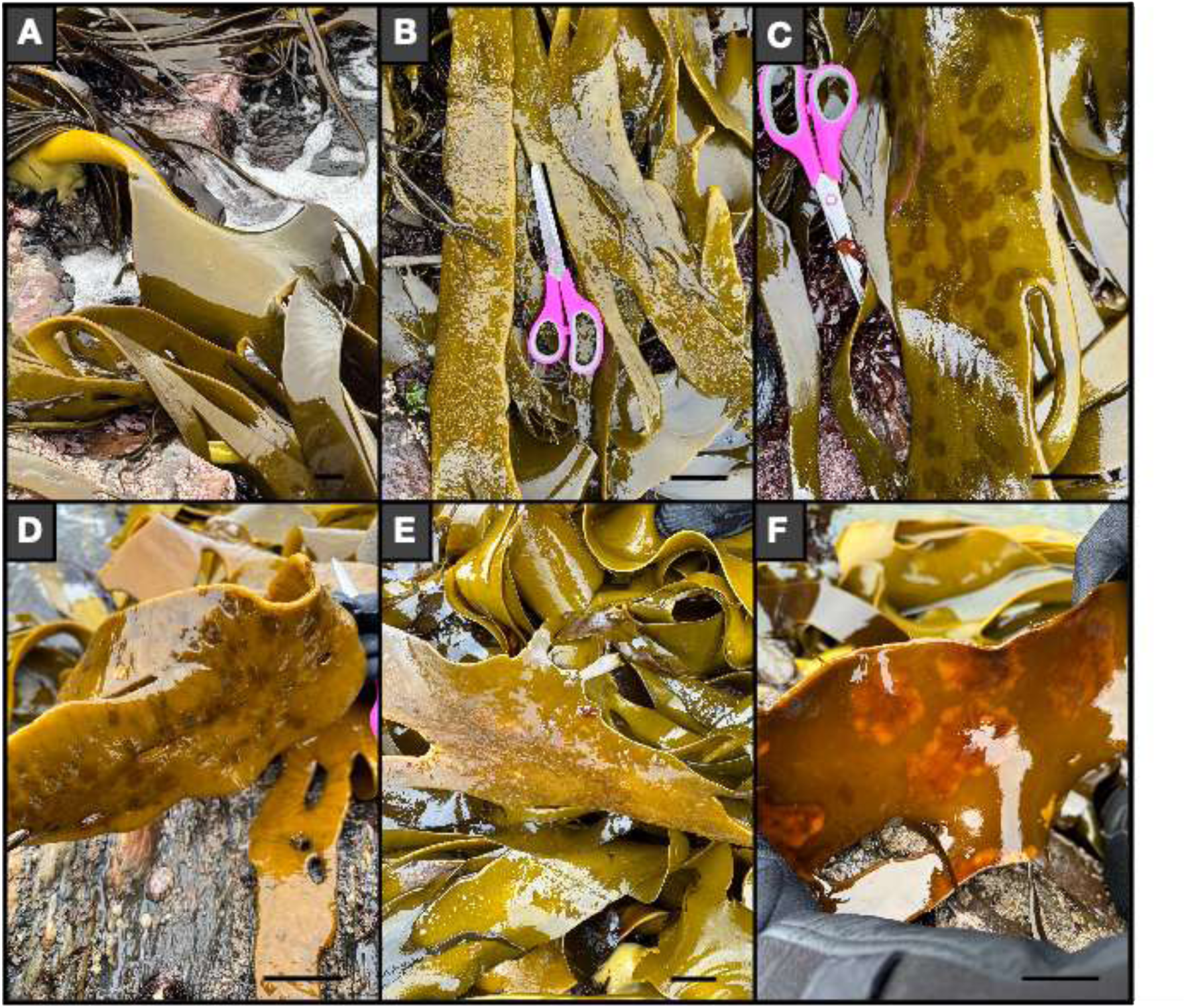
Photos of A) healthy-looking *Durvillaea* and B-F) unhealthy-looking *Durvillaea* sampled during this study. Scale bars represent ∼5 cm.

Samples were placed in a cooler during transport from sites and transferred within two hours of collection into a -80°C freezer to reduce RNA degradation, with the exception of samples collected from Tautuku which, for logistical reasons, were held at -20 °C for two days prior to transfer to -80 °C.

During the sampling period, conditions were sometimes unsuitable with high swells and wind. Due to challenging conditions, Brighton Beach was sampled twice; different areas were selected to eliminate the risk of unknowingly resampling the same individuals.

### Total RNA Extraction from Kelp Samples

Frozen kelp samples were defrosted and eight pieces (<1 mm^3^) of each were combined with 750 µL of DNA/RNA shield (ZR) in ZR BashingBead Lysis tubes (0.1 – 0.5 mm). Samples were transferred into the lysis tubes and placed into a Mini-Beadbeater-24 disrupter (Biospec Products Inc.) to be homogenised for 5 x 1-minute intervals with a 30 – 40 second cooling period, on ice, in between each homogenisation cycle to reduce RNA degradation. Total RNA was extracted using the *Quick*-RNA Plant Miniprep kits (ZR), with some additions to the manufacturer’s protocol. Specifically, Sigma-Aldrich^®^ molecular grade, pure ethanol (EtOH) washes were undertaken to remove any residual guanidine contamination. DNase I treatment was performed to digest DNA and leave intact RNA. RNA (ng/μl) and purity (A_260_/A_280_) were quantified using a NanoDrop™ One/One Microvolume UV-Vis Spectrophotometer.

### RNA Pooling and Sequencing

Due to the range of symptoms observed across sites and species, as well as project resources, it was not feasible to pool samples of specific disease phenotypes separately. Instead, for each species at each site, samples were pooled into two categories: apparently unhealthy and apparently healthy tissue, yielding 18 RNA sequencing libraries (Supplementary Table 3). To generate each library, equal volumes of RNA from individual samples were combined, with total library volumes adjusted to 80–120 μl. This approach ensured that all individuals contributed material to the pool, although it does not normalise for differences in RNA yield between samples. Where more than 20 individuals were available, a subset was selected based on RNA quality and representation of symptom variation.

When sampling kelp, *Durvillaea poha* and *Durvillaea antarctica* occasionally displayed similar morphology, likely due to environmental conditions influencing phenotypic plasticity (Fraser *et al*., 2012). As a result, some individuals of *D. poha* and *D. antarctica* were classified with uncertainty. These uncertain samples were grouped with *D. antarctica*, as a larger proportion of uncertain specimens resembled *D. antarctica* and there were fewer samples for this species.

Libraries then underwent total RNA sequencing at the Australian Genome Research Facility (ACRF). Libraries were prepared using the Illumina Stranded Total RNA Prep with Ribo-Zero Plus (Illumina) and 16 cycles of PCR. Paired-end 150 bp sequencing was performed on the Illumina NovaSeqX platform.

### Virus Transcript Assembly, Identification and Abundance

Paired-end reads were trimmed using Trimmomatic (Bolger *et al*., 2014) and their quality was assured using FastQC (Andrews, 2010). Reads were then assembled *de novo* using megahit v.1.2.9 (Li *et al*., 2015). Sequence similarity searches were performed on the assembled transcripts against a local copy of NCBI’s non-redundant (nr) protein database using Diamond (BLASTx; Buchfink *et al*., 2021). Contigs were categorised into higher kingdoms (i.e. eukaryotes, bacteria, archaea or viruses) using the Diamond (BLASTx) ‘sskingdoms’ flag option. Viral assignments were then refined by sorting BLASTx results by percentage identity and retaining only contigs with significant similarity to viral proteins (e-value ≤ 1e^−10^). For each contig, top hits were checked to confirm consistency of taxonomic assignment (e.g., multiple hits within the same family such as Narnaviridae). Contigs with ambiguous matches or without confident similarity to known viral proteins were retained as “unclassified.” Non-viral blast hits including host contigs with sequence similarity to viral transcripts (e.g. endogenous viral elements) were removed from further analysis during manual screening. Based on the Diamond results, putative viral contigs were further analysed using Geneious Prime (v 2022.2.2) to find and translate open reading frames. Viral abundances were estimated using Bowtie2 (v.2.4.4; Langmead & Salzberg, 2012), where reads were mapped onto the assembled transcripts. Similar to previous work (e.g. Waller *et al*., 2024), viral transcripts with expected abundances of less than 0.1% of the highest expected abundance for that virus across other libraries were removed from further analysis due to the possibility of index-hopping.

### Phylogenetic Analysis of Kelp Viruses

The methodology for viral phylogeny analysis was similar to Waller *et al*. (2024). As total RNA was sequenced, phylogenetic analyses focused on conserved regions and were restricted to contigs encoding RNA-dependent RNA polymerase (RdRp). Translated RdRp sequences used for phylogenetic reconstruction ranged from 78 to 1098 amino acid residues, reflecting variation in contig length across viral lineages. Translated RdRp sequences were aligned with representative protein sequences from the same virus family or other taxonomic group, acquired from NCBI RefSeq, along with the closest BLASTp hit, using MAFFT v7.490 (Katoh *et al*., 2002). Top genetic matches were selected by the lowest e-value, as some with a high percentage identity had a low query cover. Ambiguously aligned regions were removed with trimAL v1.2rev59, using a gap threshold set to 0.9 (Capella-Gutiérrez *et al*., 2009). Maximum likelihood phylogenetic trees for each virus family/group were estimated using IQ-TREE v1.6.12 (Nguyen *et al*., 2015). The LG amino acid substitution model was applied with 1000 ultra-fast bootstrapping replicates, and Figtree v1.4.4 was used to illustrate phylogenetic trees (Rambaut, 2017).

A virus was provisionally classified as a novel species if it shared less than 90% amino acid similarity in the most conserved region (i.e., RdRp/polymerase; Wang *et al*., 2017; King et al., 2012). A provisional virus (common) name has been given to novel virus sequences before formal verification by the International Committee on Taxonomy of Viruses (ICTV; Lefkowitz *et al*., 2018). Only RdRp-containing contigs were formally assessed for novelty; contigs without RdRp were classified by BLASTx similarity where possible or retained as unclassified. Viruses were termed ‘kelp-associated’ if they were closely related to previously described environmental-related viruses (such as marine viruses); in contrast viruses were termed ‘kelp viruses’ if they were closely related to previously described algal/seaweed viruses.

### Exploring Kelp Virus Richness

The library for sample LWD was removed from further analysis as no viruses were identified. Plots and analyses were completed in RStudio v4.1.0. Alpha diversity, representing the diversity or richness of species observed at a defined local scale (Andermann *et al*., 2022), was visualized using plots generated with the R package ggplot2 v3.4.1 (Wickham, 2016). Viral abundance (RPM) was analysed in relation to health status, site, and their interaction using ANOVAs, while richness was analysed with a GLM using a Poisson link, as it represents count data. Tukey HSD post-hoc tests were used for further analysis of viral abundance.

## Results

One library (healthy *D. poha* from Lawyer’s Head) was unable to be sequenced as it did not pass AGRF quality controls checks (Supplementary Table 4). The 17 other libraries succeeded, however, the library containing unhealthy *D. willana* samples from Lawyer’s Head returned no viral sequences so was removed from the analyses to reduce bias in site comparisons. The number of raw viral reads for each library ranged from 732,000 – 227 million (Supplementary Table 4). The percentage of viral reads ranged from 0% to 0.044% (Supplementary Table 4).

### Viral Abundance and Richness

Viral abundance was quantified using reads per million (RPM), a metric that normalises sequencing depth between samples by dividing the total number of reads in each sample by one million. The abundance of viral reads was significantly different across sites (ANOVA, *F*_2,10_ = 4.97, *p* = 0.032) and the interaction between health status and site (ANOVA, *F*_2,10_ = 4.88, *p* = 0.033) but was not significantly different in terms of health status (ANOVA, *F*_1,10_ = 0.51, *p* = 0.49) (Fig. 3A). Tukey multiple pairwise comparison of means show the abundance of viral reads was significantly higher at Lawyer’s Head compared to Brighton Beach with a mean difference of 12,482 RPM (95% CI: 1,369.3 to 23,595, *p* = 0.029). The abundance of viral reads was significantly higher in unhealthy Lawyer’s Head samples compared to unhealthy Brighton Beach samples (*p* = 0.043), healthy Brighton Beach samples (*p* = 0.027) and unhealthy Tautuku Peninsula samples (*p* = 0.026).

**Fig. 3.**
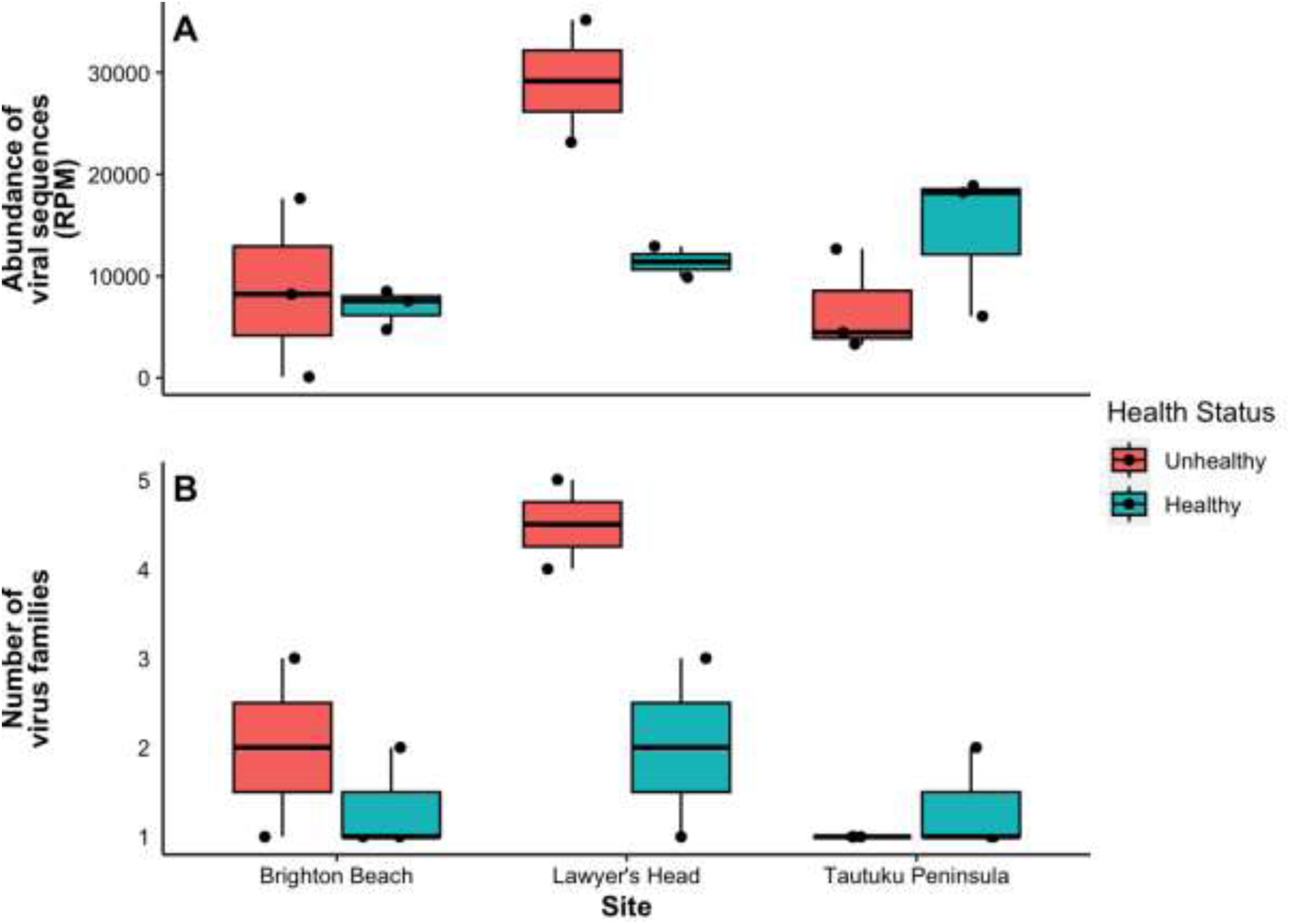
Boxplots showing (A) abundance of viral reads (RPM) and (B) virus family richness in *Durvillaea* samples by site and health status. Boxes indicate the median and interquartile range, with whiskers extending to 1.5× the interquartile range. Points represent individual biological replicates. Brighton Beach and Tautuku Peninsula (n = 3 each) and Lawyer’s Head (n = 2).

A GLM showed virus family richness did not significantly differ for healthy samples compared to unhealthy samples (Estimate = -0.41, SE = 0.65, z = -0.628, *p* = 0.090) or across sites (Estimate = -0.4055, SE = 0.6455, z = -0.628, *p* = 0.090) (Fig. 3B). Virus family richness at Lawyer’s Head was not significantly different to Brighton Beach (Estimate = 0.81, SE = 0.53, z = 1.54, *p* = 0.12) and richness at Tautuku Peninsula was not significantly different to Brighton Beach (Estimate = -0.69, SE = 0.71, z = -0.980, *p* = 0.33) (Fig. 3B).

Across virus families, read abundance tended to be higher in unhealthy kelp, with the exception of *Narnaviridae* and Ormycovirus, which showed similar RPM in healthy and unhealthy tissue (Fig. 4; Fig. 5 & Supplementary Table 4). Statistical comparisons at the family level were not possible due to pooling of samples. Viral abundance data (reads per million; RPM) was log₁₀-transformed [log₁₀(RPM + 1)] for visual comparison with virus family and health status to account for zero inflation and right-skewed distributions. This transformation improves comparability across virus families spanning multiple orders of magnitude in abundance.

**Fig. 4.**
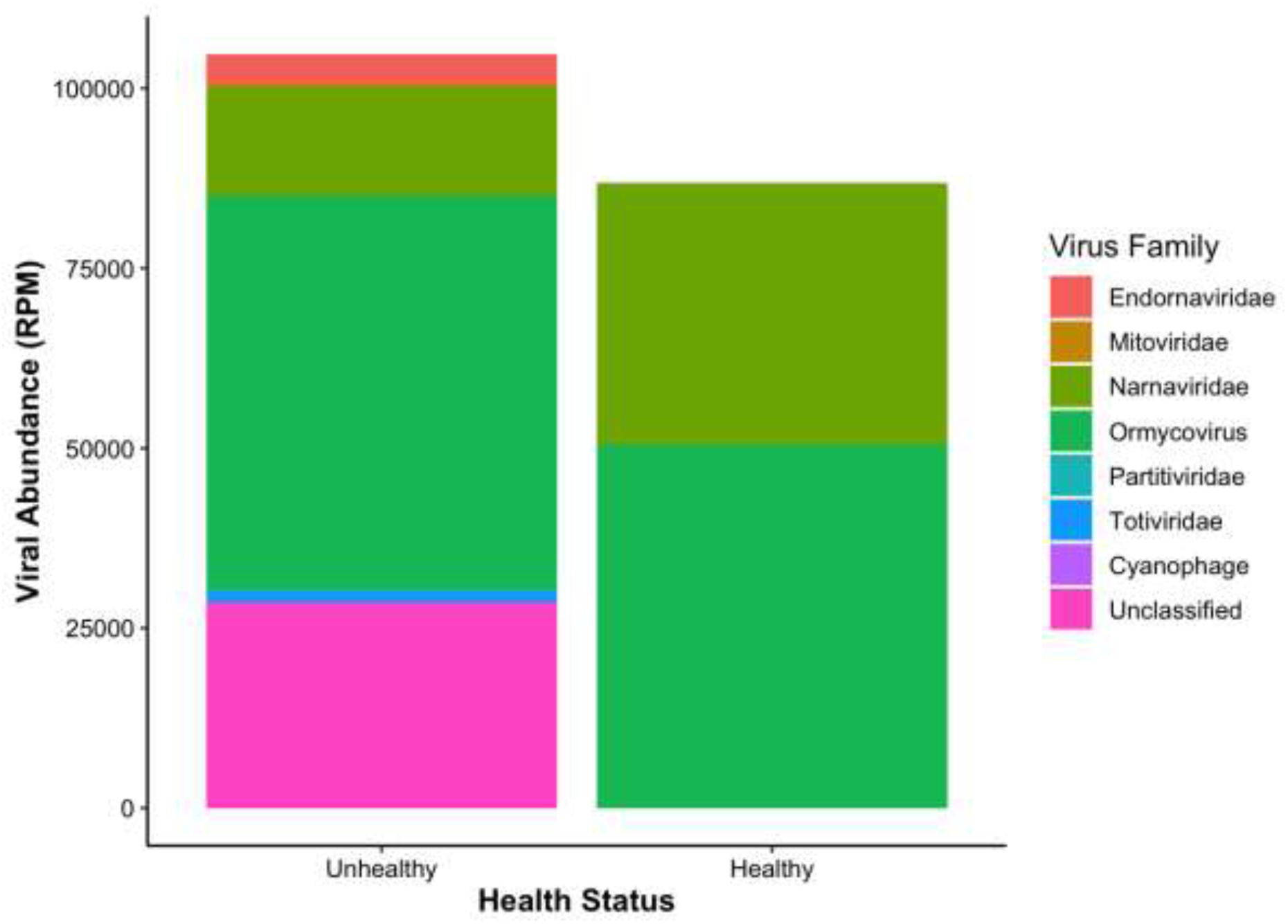
Composition of viral abundance across virus families in healthy and unhealthy *Durvillaea*. Bars represent total reads per million (RPM) summed across all samples within each health status (n = 8 per group) and stacked by virus family.

**Fig. 5.**
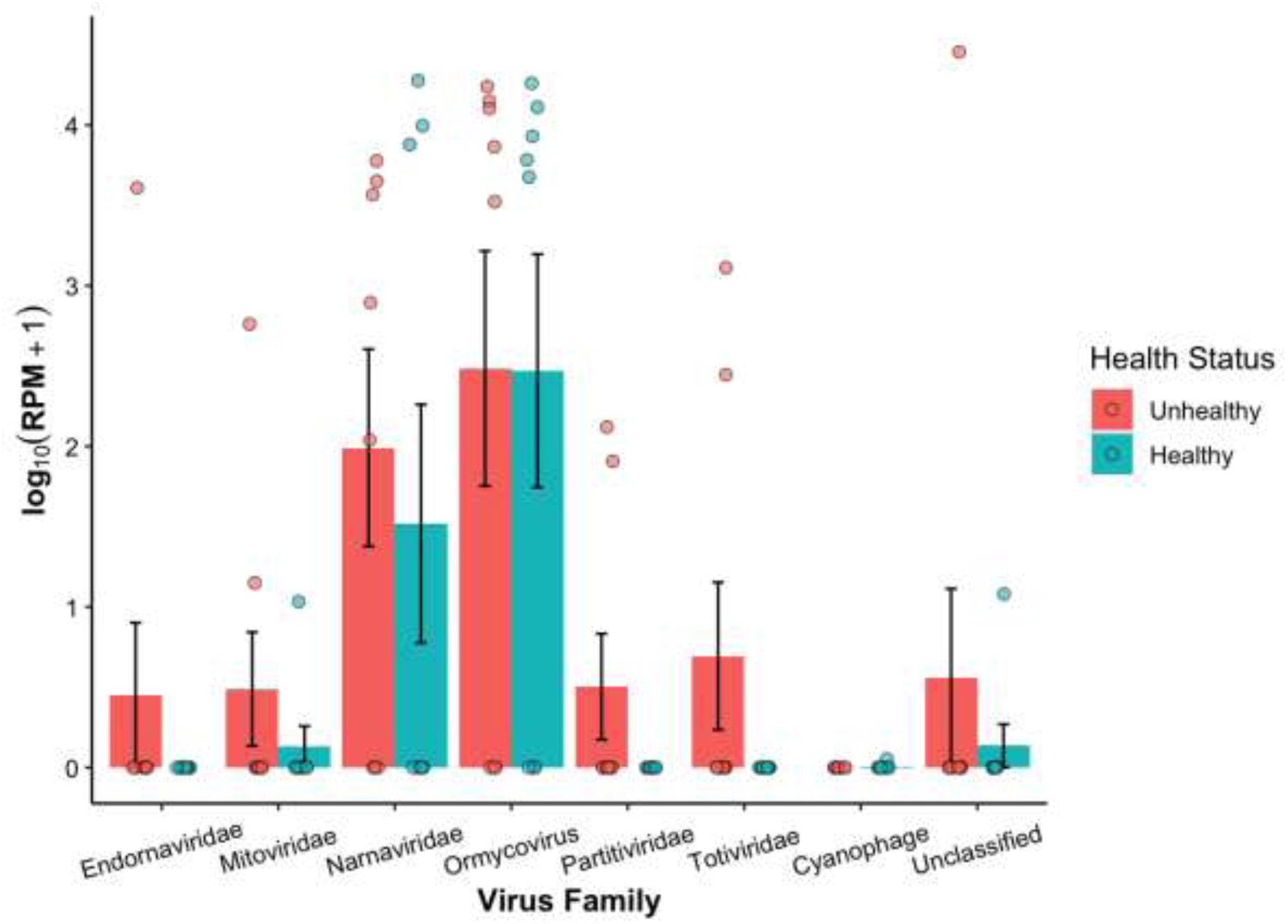
Viral abundance across virus families in healthy and unhealthy *Durvillaea* samples. Abundance is shown as log₁₀-transformed reads per million (RPM + 1). Bars represent group means, error bars represent ±1 standard error of the mean (SE), and points represent individual biological replicates (n = 8 per group).

### Virus Family Phylogeny

Phylogenetic analysis of RdRp sequences revealed five virus families/groups in *Durvillaea*: *Mitoviridae*, *Narnaviridae*, *Totiviridae*, *Partitiviridae*, and Ormycovirus. Many sequences clustered closely in the trees, likely reflecting the use of conserved regions for tree reconstruction rather than complete genome identity. Amino acid coverage for classification varied from 30–75% (Supplementary Table 6), which may obscure Across virus families, read abundance appeared higher in unhealthy kelp for most families, with the exception of Narnaviridae and Ormycoviridae, which showed similar RPM in healthy and unhealthy tissue (Supplementary Table 4). Statistical comparisons at the individual family level were not possible due to pooling of samples, so these observations are descriptive. in less-conserved regions. Most sequences were detected in both healthy and unhealthy kelp, and no lineages were found exclusively in healthy tissue, although pooling and coverage limitations prevent definitive conclusions about health-specific virus subtypes.

### Mitoviridae

Four mitoviruses were identified in unhealthy *Durvillaea antarctica* from Lawyer’s Head and one mitovirus was identified in healthy *Durvillaea willana* from Lawyer’s Head. Kelp-associated mitovirus 1 (GenBank accessions PV404153.1 and XQW56660.1) sequenced from unhealthy *D. antartica* at Lawyer’s Head shared 46.53% amino acid sequence similarity to *Proteus mito-like virus* (DAZ87243.1) which was previously identified in the chlorophyte *Polyblepharides amylifera* (Fig. 6 & Supplementary Table 5; Charon *et al*., 2021). Kelp-associated mitovirus 2 (GenBank accessions PV404154.1 and XQW56661.1) sequenced from unhealthy *D. antartica* at Lawyer’s Head shared 44.59% amino acid sequence similarity to *Ripatrop virus* (WZH61242.1) which was previously identified in riverbank sediment from Australia (Fig. 6 & Supplementary Table 5; Sadiq *et al*., 2024). Kelp mitovirus 1 (GenBank accessions PV404155.1 and XQW56662.1) sequenced from unhealthy *D. antartica* at Lawyer’s Head shared 58.62% amino acid sequence similarity to two *Mitovirus sp*. (BDC79612.1 & BDC79614.1) which were previously identified from the red macroalgae *Neopyropia yezoensis* and its nori sheet products (Fig. 6 & Supplementary Table 5; Mizutani *et al*., 2022). Kelp mitovirus 2 (GenBank accessions PV404156.1 and XQW56663.1) sequenced from unhealthy *D. antartica* at Lawyer’s Head also shared 50.00% amino acid similarity to the red macroalgae Mitovirus sp. (BDC79612.1) (Fig. 6 & Supplementary Table 5; Mizutani *et al*., 2022). Kelp-associated mitovirus 3 (GenBank accessions PV404157.1 and XQW56664.1) sequenced from healthy *D. willana* at Lawyer’s Head shared 45.59% amino acid sequence similarity to *Mitoviridae sp.* (URG16825.1) which was previously identified in sheep faeces from China (Fig. 6 & Supplementary Table 5; Chen *et al*., 2022).

**Fig. 6.**
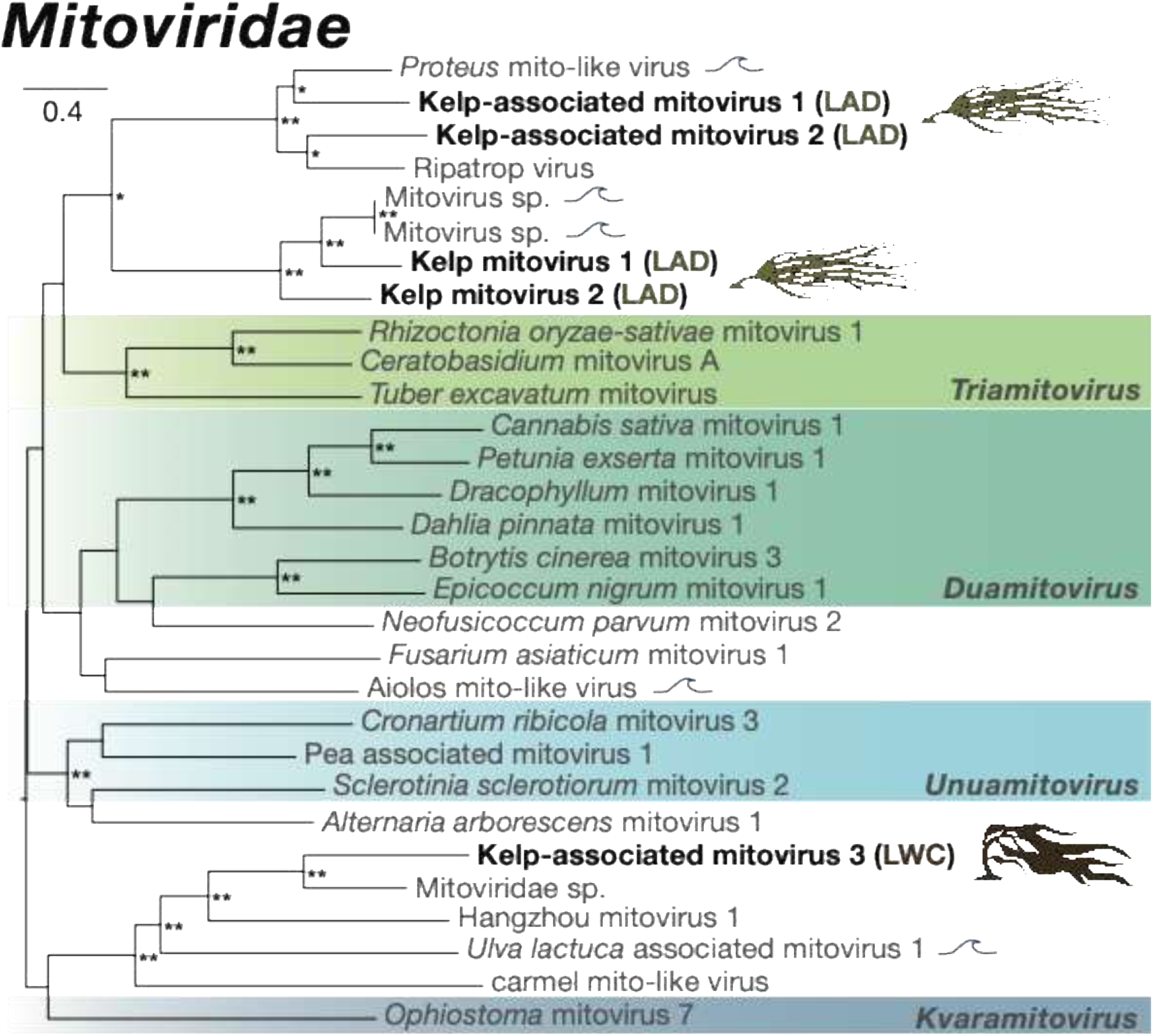
Maximum likelihood phylogenetic tree of representative viral transcripts from the family *Mitoviridae*. *Durvillaea* RdRp viral transcripts sequenced during this study are shown in bold. Viral sequences were obtained from RNA sequencing data collected across three different sites and from three species of *Durvillaea*. Known genera have been coloured. The phylogenetic tree was mid-point rooted for clarity and nodes with bootstrap values of 75 – 89 are noted with a single asterisk and bootstrap values of 90 – 100 are noted with a double asterisk. The branches are scaled according to the number of amino acid substitutions per site. The wave represents viruses sequenced from the marine environment.

### Narnaviridae

Four narnaviruses were identified in unhealthy *Durvillaea spp.*, and one narnavirus was identified in healthy *D. antarctica*. Kelp-associated narnavirus 1 was identified in healthy *D. antartica* at Brighton Beach (GenBank accessions PV404162.1 and XQW56669.1); Lawyer’s Head (GenBank accessions PV404164.1 and XQW56671.1); and Tautuku Peninsula (GenBank accessions PV404163.1 and XQW56670.1), unhealthy *D. antartica* at Brighton Beach (GenBank accessions PV404158.1 and XQW56665.1); Lawyer’s Head (GenBank accessions PV404159.1 and XQW56666.1); and Tautuku Peninsula (GenBank accessions PV404161.1 and XQW56668.1), and unhealthy *D. poha* at Lawyer’s Head (GenBank accessions PV404160.1 and XQW56667.1) and shared 48.12 - 61.78% amino acid sequence similarity to *Plasmopara viticola lesion associated narnavirus 10* (QIR30289.1) which was previously identified in the oomycete (*Plasmopara viticola*) that is the causal agent of downy mildew (Fig. 7 & Supplementary Table 5; Chiapello *et al*., 2020). Kelp-associated narnavirus 2 was sequenced from unhealthy *D. poha* libraries at Brighton Beach (GenBank accessions PV404165.1 and XQW56672.1) and Lawyer’s Head (GenBank accessions PV404166.1 and XQW56673.1), and shared 47.27% and 45.24% amino acid sequence similarity respectively *Blattella germanica narna-like virus 1* (DBA56653.1) which was previously identified in German cockroaches (*Blattella germanica*) from USA and Spain (Fig. 7 & Supplementary Table 5; Wu *et al*., 2024). Kelp-associated narnavirus 3 (GenBank accessions PV404167.1 and XQW56674.1) was sequenced from unhealthy *D. antarctica* at Brighton Beach and also shared 43.26% amino acid sequence similarity to the same *Blattella germanica narna-like virus 1* (DBA56653.1; Fig. 7 & Supplementary Table 5; Wu *et al*., 2024). Kelp-associated narnavirus 4 (GenBank accessions PV404168.1 and XQW56675.1) was sequenced from unhealthy *D. antarctica* at Lawyer’s Head and shared 61.68% amino acid similarity to *Narnaviridae sp.* (UJQ92711.1) which was previously identified from soil samples in China (Fig. 7 & Supplementary Table 5; Chen *et al*., 2022).

**Fig. 7.**
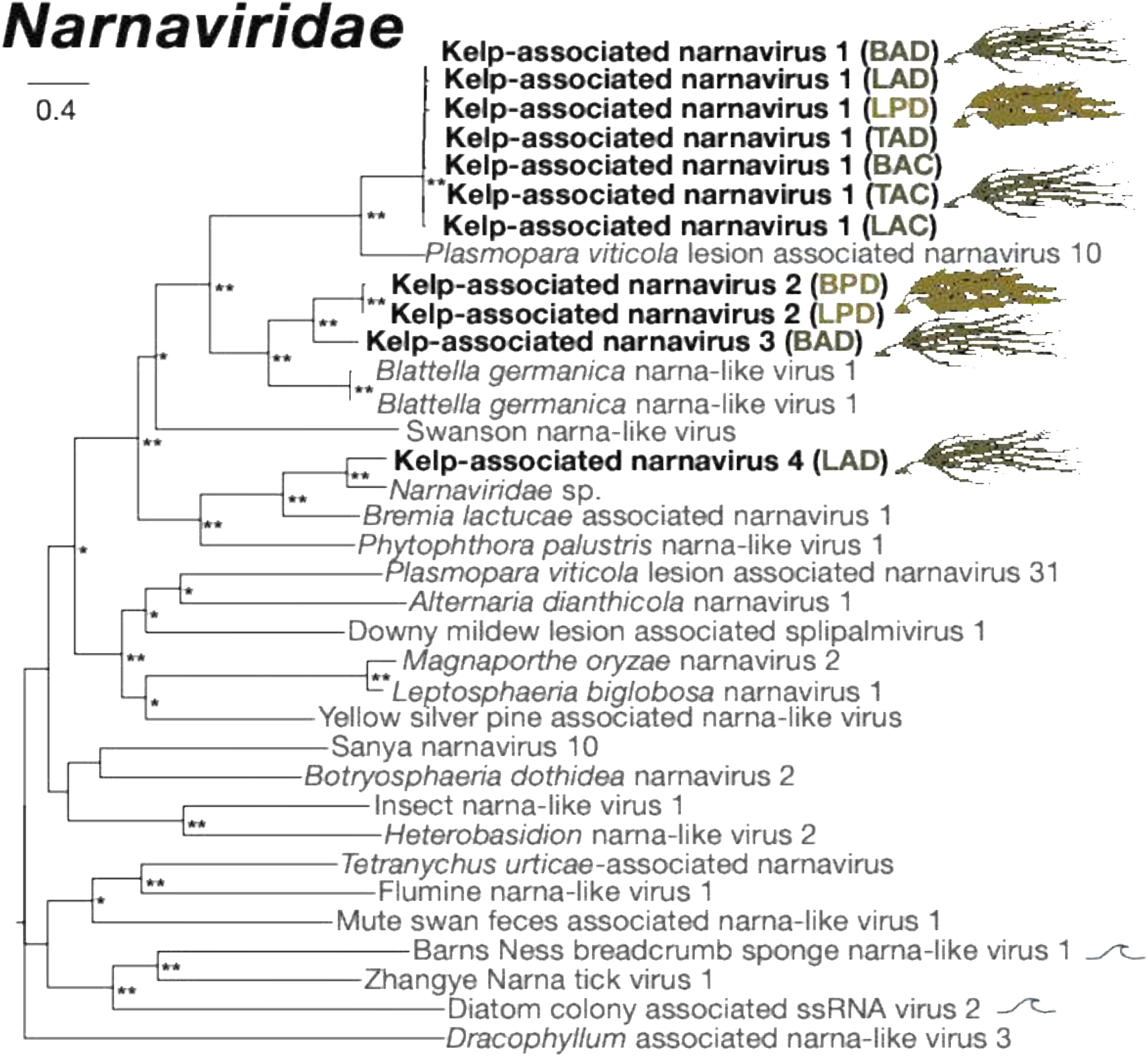
Maximum likelihood phylogenetic tree of representative viral transcripts from the family *Narnaviridae*. *Durvillaea* RdRp viral transcripts sequenced during this study are shown in bold. Viral sequences were obtained from RNA sequencing data collected across three different sites and from three species of *Durvillaea*. The phylogenetic tree was mid-point rooted for clarity and nodes with bootstrap values of 75 – 89 are noted with a single asterisk and bootstrap values of 90 – 100 are noted with a double asterisk. The branches are scaled according to the number of amino acid substitutions per site. The wave represents viruses sequenced from the marine environment.

### Ormycovirus

Two ormycoviruses were identified in unhealthy *Durvillaea poha*, and one ormycovirus was identified in healthy *Durvillaea poha*. Kelp-associated ormycovirus 1 was sequenced from healthy *D. poha* at Brighton Beach (GenBank accessions PV404172.1 and XQW56679.1); and Tautuku Peninsula (GenBank accessions PV404169.1 and XQW56676.1), and unhealthy *D. poha* at Brighton Beach (GenBank accessions PV404173.1 and XQW56680.1); Lawyer’s Head (GenBank accessions PV404171.1 and XQW56678.1); and Tautuku Peninsula (GenBank accessions PV404170.1 and XQW56677.1) and shared 29.18 - 37.28% amino acid sequence similarity to *Phytophthora cinnamomi ormycovirus 9-16* (XDO01627.1) which was previously identified from a soil-bourne water mould commonly known as cinnamon fungus (*Phytophthora cinnamomi*) in Japan (Fig. 8 & Supplementary Table 5). Kelp-associated ormycovirus 2 was sequenced from unhealthy *D. poha* at Brighton Beach (GenBank accessions PV404174.1 and XQW56681.1); and Lawyer’s Head (GenBank accessions PV404175.1 and XQW56682.1) and also shared 43.29% and 33.84% amino acid sequence similarity respectively with the same *Phytophthora cinnamomi ormycovirus 9-16* (XDO01627.1) (Fig. 8 & Supplementary Table 5).

**Fig. 8.**
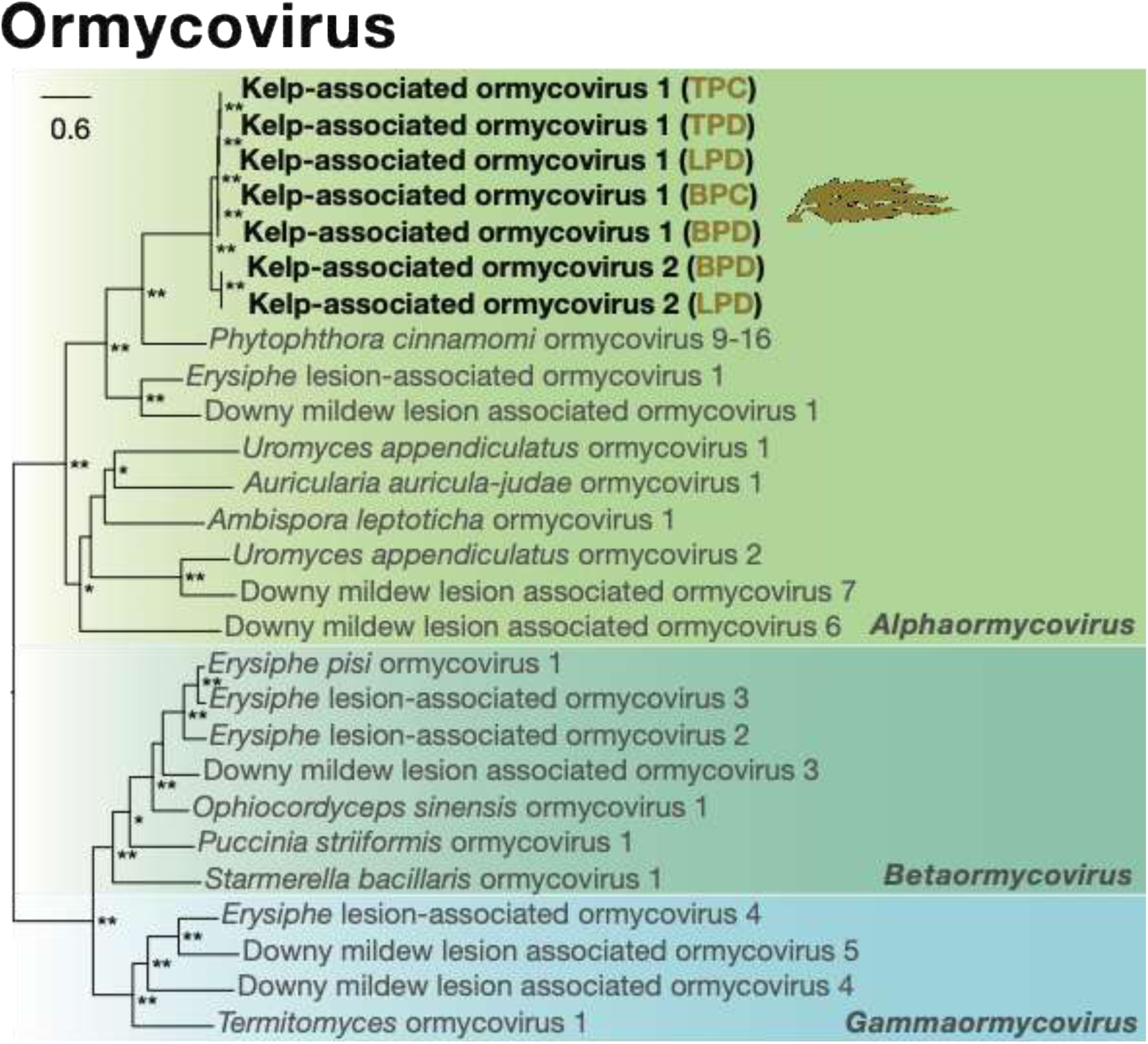
Maximum likelihood phylogenetic tree of representative viral transcripts from the group Ormycovirus. *Durvillaea* RdRp viral transcripts sequenced during this study are shown in bold. Proposed groups within the classification Ormycovirus have been coloured. Viral sequences were obtained from RNA sequencing data collected across three different sites and from three species of *Durvillaea*. The phylogenetic tree was mid-point rooted for clarity and nodes with bootstrap values of 75 – 89 are noted with a single asterisk and bootstrap values of 90 – 100 are noted with a double asterisk. The branches are scaled according to the number of amino acid substitutions per site. The wave represents viruses sequenced from the marine environment.

### Partitiviridae

Three partitiviruses were identified in unhealthy *Durvillaea* spp.. Kelp-associated partitivirus 1 (GenBank accessions PV404176.1 and XQW56683.1) was sequenced from unhealthy *D. antarctica* at Lawyer’s Head and shared 74.26% amino acid sequence similarity to *Partitiviridae sp.* (UDL14432.1) which was previously identified in lake sediment from China (Fig. 9 & Supplementary Table 5; Chen *et al*., 2022). Kelp-associated partitivirus 2 (GenBank accessions PV404177.1 and XQW56684.1) was sequenced from unhealthy *D. poha* at Brighton Beach and shared 33.94% amino acid sequence similarity to *Aegean partiti-like virus* (DAZ87271.1) which was previously identified in marine chlorophyte (*Tetraselmis chuii*) (Fig. 9 & Supplementary Table 5; Charon *et al*., 2021). Kelp-associated partitivirus 3 (GenBank accessions PV404178.1 and XQW56685.1) was sequenced from unhealthy *D. antarctica* at Lawyer’s Head and shared 37.12% amino acid sequence similarity to *Diatom colony associated dsRNA virus 2* (YP_009551448.1) which were previously identified from a diatom colony on tidal rocks from Japan (Fig. 9 & Supplementary Table 5; Urayama *et al*., 2016).

**Fig. 9.**
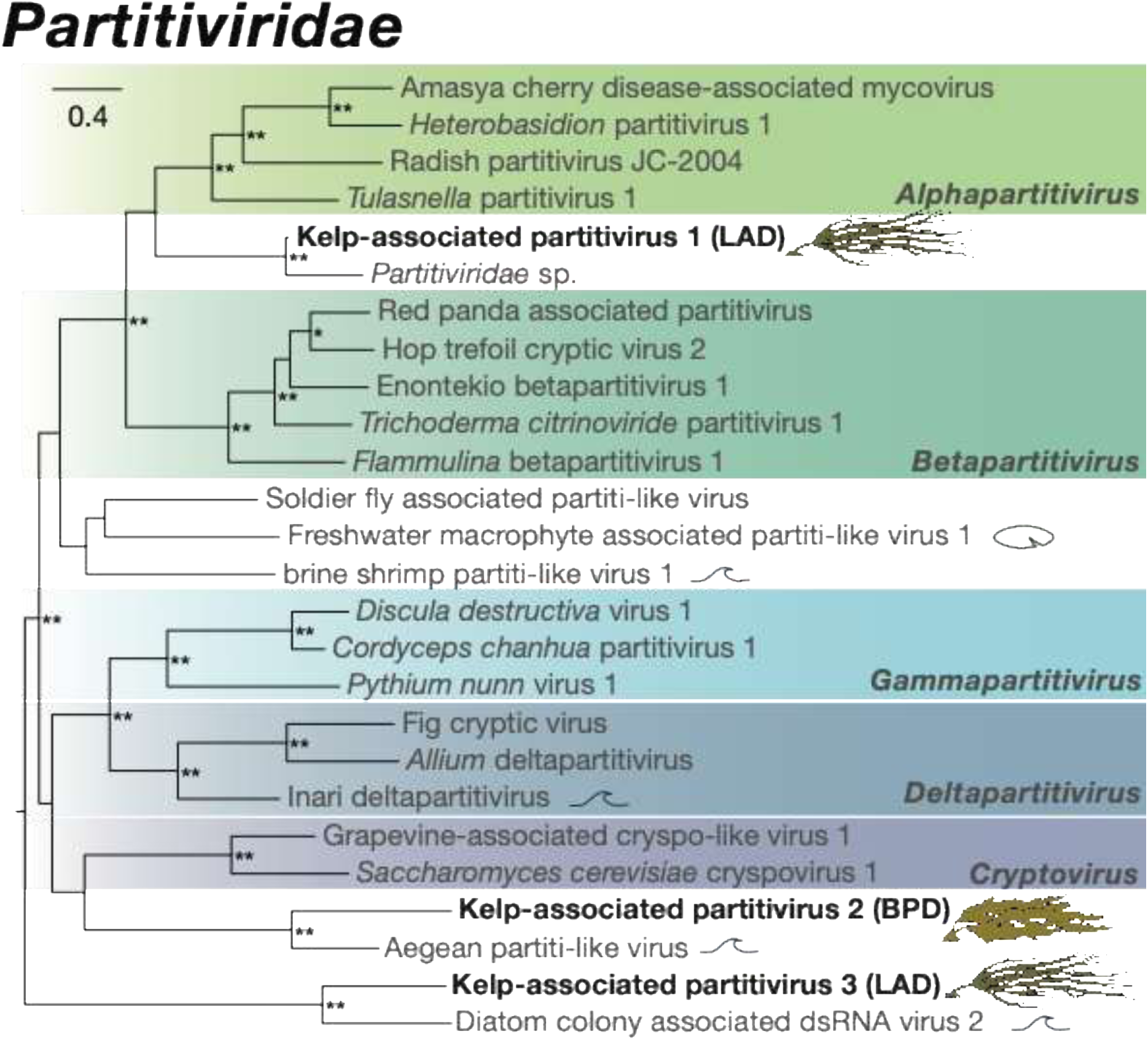
Maximum likelihood phylogenetic tree of representative viral transcripts from the family *Partiviridae*. *Durvillaea* RdRp viral transcripts sequenced during this study are shown in bold. Known genera have been coloured. Viral sequences were obtained from RNA sequencing data collected across three different sites and from three species of *Durvillaea*. The phylogenetic tree was mid-point rooted for clarity and nodes with bootstrap values of 75 – 89 are noted with a single asterisk and bootstrap values of 90 – 100 are noted with a double asterisk. The branches are scaled according to the number of amino acid substitutions per site. The wave represents viruses sequenced from the marine environment. The lily pad represents viruses sequenced from freshwater macrophytes.

### Totiviridae

Four viruses from the family *Totiviridae* were identified in unhealthy *Durvillaea* spp.. Kelp-associated toti-like virus 1 (GenBank accessions PV404179.1 and XQW56686.1) was sequenced from unhealthy *D. poha* at Lawyer’s Head and shared 60.34% amino acid sequence similarity to *Diatom totivirus 1* (BBZ90080.1) which was previously identified in a diatom (*Melosira* sp.) from Japan (Fig. 10 & Supplementary Table 5; Chiba *et al*., 2020). Kelp-associated toti-like virus 2 (GenBank accessions PV404180.1 and XQW56687.1) was sequenced from unhealthy *D. poha* at Lawyer’s Head and also shared 44.62% amino acid sequence similarity to the same *Diatom totivirus 1* (BBZ90080.1) (Fig. 10 & Supplementary Table 5; Chiba *et al*., 2020). Kelp-associated toti-like virus 3 (GenBank accessions PV404181.1 and XQW56688.1) was sequenced from unhealthy *D. poha* at Lawyer’s Head and shared 54.29% amino acid sequence similarity to *Diatom colony associated virus-Like RNA Segment 4* (BAU79524.1) which were previously identified from a diatom colony on tidal rocks from Japan (Fig. 10 & Supplementary Table 5; Urayama *et al*., 2016). Kelp-associated toti-like virus 4 (GenBank accessions PV404182.1 and XQW56689.1) was sequenced from unhealthy *D. willana* at Brighton Beach and shared 39.87% amino acid sequence similarity to *Phytophthora palustris toti-like virus 5-2* (WAK73616.1) which were previously identified from a plant-damaging oomycete (*Phytophthora palustris*) from Indonesia (Fig. 10 & Supplementary Table 5; Botella *et al*., 2022).

**Fig. 10.**
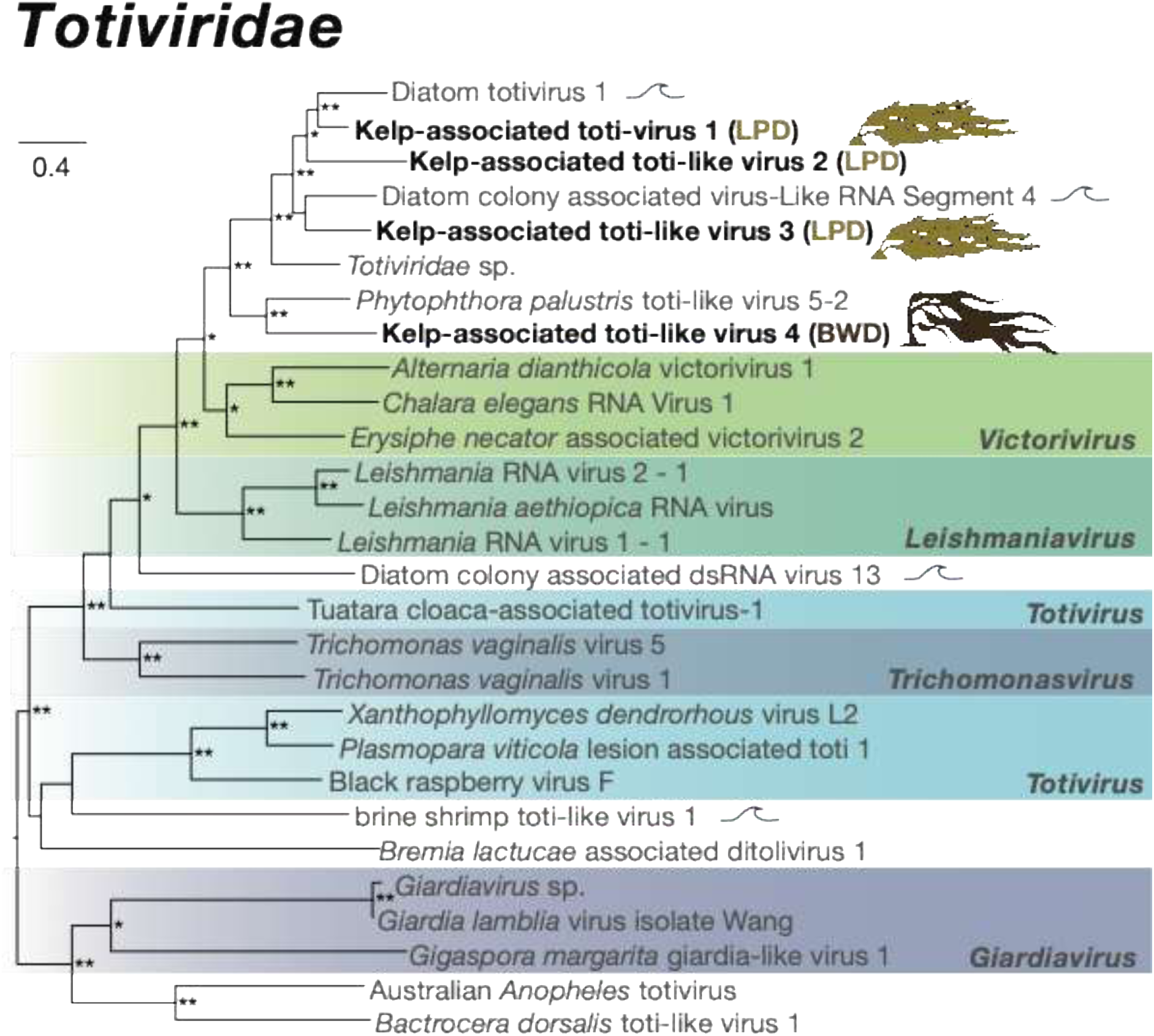
Maximum likelihood phylogenetic tree of representative viral transcripts from the family *Totiviridae*. *Durvillaea* RdRp viral transcripts sequenced during this study are shown in bold. Known genera have been coloured. Viral sequences were obtained from RNA sequencing data collected across three different sites and from three species of *Durvillaea*. The phylogenetic tree was mid-point rooted for clarity and nodes with bootstrap values of 75 – 89 are noted with a single asterisk and bootstrap values of 90 – 100 are noted with a double asterisk. The branches are scaled according to the number of amino acid substitutions per site. The wave represents viruses sequenced from the marine environment.

## Discussion

Our hypothesis that we would discover diverse viruses was supported; indeed, all viruses sequenced in this study were novel, with RdRp viral transcripts clustering within four virus families—*Mitoviridae*, *Narnaviridae*, *Totiviridae*, and *Partitiviridae*—along with one recently proposed group, Ormycovirus. The abundance of viral sequences and number of virus families was not significantly different between healthy and unhealthy kelp samples. RdRp Totivirus and partitivirus sequences were only identified in unhealthy samples, suggesting a potential association, which could be explored in future studies. Therefore, the hypothesis that apparently unhealthy tissue would yield different viral sequences compared to ‘healthy’ tissue was only partially supported. Furthermore, the detection of virus families typically associated with fungi and other organisms raises questions about potential cross-kingdom viral transmission and host specificity, requiring further investigation into the ecological roles and pathogenic potential of these viruses.

Among the viral sequences identified in *Durvillaea*, *Mitoviridae* appears to be the only family with viruses potentially directly infecting the kelp (kelp mitovirus 1 & 2; Fig. 6). Mitoviruses (family *Mitoviridae*; order *Cryppavirales*) are non-segmented, positive-sense single stranded RNA viruses that lack a capsid (Wolf *et al*., 2018; Jacquat *et al*., 2022). Mitoviruses are typically associated with fungi, but have been identified in invertebrates, plants and marine environments, replicating in the cytosol and mitochondria of hosts (Shi *et al*., 2016; Hillman & Cai, 2013; Nerva *et al*., 2019). Their detection in *Durvillaea* may indicate a novel host-virus interaction. Two novel viruses sequenced from *Durvillaea antarctica* from Lawyer’s Head were most closely related to mitoviruses sequenced from *Neopyropia yezoensis*, a species of red macroalgae, as well as processed nori sheets derived from this species (Mizutani *et al*., 2022). The authors speculated that mitoviruses were directly infecting the macroalgae, but that they could be beneficial as they were identified from all of the *N. yezoensis* individuals/products (Mizutani *et al*., 2022). Even though the closest relatives of these mitoviruses were identified in macroalgae, viral hosts are often misidentified, as the true hosts may instead be the organism’s parasites, dietary components, or environmental contaminants (Cobbin *et al*., 2021).

Mizutani *et al*. (2022) identified mitoviruses from *Neopyropia yezoensis* and inferred their host association based on phylogenetic relationships. While phylogenetic inference is a widely used and informative approach for predicting viral hosts, it is not definitive and, in the absence of experimental validation, may result in misassignment—particularly in systems where host–virus interactions are not well characterised. While the effects of these mitoviruses on kelp health are not yet understood, and *Durvillaea* cannot be conclusively identified as their true host in this study, their role could range from benign to pathogenic or symbiotic interactions. To improve the accuracy of virus–host identification, future studies could incorporate methods such as coevolutionary analyses of viral and host genomes (Cobbin *et al*., 2021; Fonseca *et al*., 2020; Charon *et al*., 2019), which examine evolutionary patterns to infer host associations. Additionally, single-cell genomics offers a powerful approach for detecting virus–host interactions at the cellular level and could be adapted for use in macroalgal systems (Castillo *et al*., 2021; Li *et al*., 2022).

In contrast to the kelp-infecting mitoviruses, the other virus families detected — *Narnaviridae*, *Totiviridae*, *Partitiviridae*, and Ormycoviruses—as well as additional *Mitoviridae* sequences, may not be specific to kelp; rather, they could be free viruses or associated with other hosts in the marine environment. Ormycovirus is a newly proposed category of viruses with three groups: alphaormycovirus, betaormycovirus and gammaormycovirus (Forgia *et al*., 2022). Ormycoviruses are found to infect fungi and potentially plants, with further analysis required to confirm this association (Niu *et al*., 2024). Further novel viral sequences classified as ‘environmental’ were most closely related to organisms other than macroalgae. Instead, these viruses possibly originate from the surrounding marine environment, potentially linked to fungi, protists, or other microorganisms within the ecosystem, interacting indirectly with kelp. While the direct impact of environmental viruses on *Durvillaea* remains unclear, the presence of these viruses raises intriguing questions. For instance, they may contribute to the broader marine virome and indirectly affect kelp by interacting with symbiotic or pathogenic microorganisms in the kelp’s microbiome (Zhao *et al*., 2024). Additionally, viral interactions with other marine species could influence nutrient cycling, food web dynamics and ecosystem processes, potentially altering kelp and ecosystem functioning (Suttle, 2007; Wilhelm & Suttle, 1999). Understanding these environmental viruses’ ecological roles could provide deeper insights into marine ecosystem complexity and stability.

The elevated viral abundance observed in *Durvillaea* samples from Lawyer’s Head, particularly in apparently unhealthy tissue, raises the possibility that local anthropogenic pressures may influence viral load in intertidal kelp. Lawyer’s Head is located near treated wastewater discharge points servicing the city of Dunedin, as well as emergency overflow outfalls that operate during heavy rainfall events (Dunedin City Council, 2016). Wastewater effluent is known to contain high numbers of microorganisms and can alter coastal environments through nutrient enrichment, contaminants, and changes to abiotic conditions, all of which may influence host–pathogen interactions and disease dynamics (Bojko et al., 2020). While the present study cannot establish a causal link between anthropogenic inputs and viral abundance due to the absence of environmental measurements, this interpretation is supported by a recent regional survey of the same kelp genus that reported increased disease-like phenotypes closer to sewage outlets along the Otago coastline (de Klein et al., 2026). Future studies integrating virome analyses with environmental and pollution metrics will be critical for determining how human activities shape viral dynamics in macroalgal hosts.

A significant limitation of this study arises from pooling samples by site, species, and health status, which hinders the ability to directly link specific virus types to observed symptoms in *Durvillaea*. This approach, while practical for increasing sample size and identifying broader viral dynamics, prevents the resolution of whether individual viral species are associated with particular disease-like symptoms or host responses. As a result, the ecological and pathogenic significance of the viruses detected remains unclear, requiring future studies that focus on individual sampling to establish more direct host-virus relationships. Another limitation was that pooling was performed by equal RNA volumes rather than normalised concentrations, meaning that individuals with higher or lower RNA yield may have been over- or under-represented in the pooled libraries. Pooling also reduced the number of independent replicates, limiting our ability to statistically detect differences in viral community composition between health states or sites. However, this approach was deemed necessary given project resources, and did ensure broad inclusion of samples across sites, species, and symptom types. Despite these limitations, the data provide intriguing insights into the virome of intertidal coastal kelp in New Zealand, and open exciting new avenues for future research.

The timing of our collections could have influenced which viruses were detected. The marine virome may vary throughout the year due to fluctuations in environmental variables, including temperature, salinity, chlorophyll *a* levels, nutrient availability, and dissolved oxygen concentrations. These seasonal changes can influence the abundance, presence and composition of viruses (Hwang *et al*., 2017). Many studies on seasonal dynamics of marine viruses have focused on viruses associated with the microbial community, including phytoplankton and bacteria (Suttle, 2007). Viral abundances associated with the microbial community have been observed to be higher in late summer or autumn, and lower in winter or early spring, likely due to coinciding peaks and declines in host bacterial abundance (Weinbauer *et al*., 1995) and chlorophyll-*a* levels (Gainer *et al*., 2017). Seasonal variations in viral abundance can differ depending on the viral species (Hevroni *et al*. 2020). The complexity of viral dynamics, coupled with the limited research on viruses infecting macroalgae, makes it challenging to assess whether autumn was an appropriate time for sampling in terms of virus detection. Further research, including longitudinal studies across multiple seasons, is necessary to determine if significant fluctuations in abundance and diversity occur.

Our research has revealed that diverse novel viruses are associated with intertidal macroalgae in New Zealand, and hints at possible anthropogenic influences on viral loads. Further research on this important topic will help us understand the pathogenicity of these viruses, and environmental factors influencing their presence and abundance; nonetheless, this first snapshot of viral diversity provides a useful foundation for addressing those critical questions.

## Data Availability

All raw metagenomic sequence data generated in this study have been deposited in the NCBI Sequence Read Archive (SRA) under BioProject accession number PRJNA1235276. The dataset comprises algal metagenome samples (Taxonomy ID: 1300146) with sample accession numbers SAMN47323595–SAMN47323612 and corresponding SRA run accessions SRR32669716–SRR32669707. The full dataset is publicly available at: https://www.ncbi.nlm.nih.gov/bioproject/PRJNA1235276.

## Funding

This work was supported by the Royal Society of New Zealand, via a Rutherford Discovery Fellowship to CIF [grant number RDF-UOO1803] and Marsden Fund [grant number MFP-20-UOO-173]. JLG is funded by a New Zealand Royal Society Rutherford Discovery Fellowship (RDF-20-UOO-007) and the Webster Family Chair in Viral Pathogenesis.

## Acknowledgements

Thank you to Ryan Woodhouse for help in the field. Thank you to Isa de Vries, Janelle Wierenga and Jess Darnley for help with lab and analytical methods. Thank you to Abby Smith for feedback on the draft introduction and methods. This research was undertaken as an Honours project by LH.

## Conflict of Interest

The authors declare no conflict of interest.

## Author Contributions

**Lily Harvey:** conceptualisation, investigation, formal analysis, writing – original draft preparation; **Stephanie Waller:** investigation, formal analysis, writing – review and editing; **Rebecca French:** investigation, formal analysis, writing – review and editing; **Jemma Geoghegan:** conceptualisation, data curation, supervision, writing – review and editing; **Ceridwen Fraser:** conceptualisation, funding acquisition, resources, supervision, writing – review and editing.

## Notes

https://www.ncbi.nlm.nih.gov/bioproject/PRJNA1235276

### Competing Interest Statement

The authors have declared no competing interest.

